# KaicoTracker: a robust and automated locomotory analysis for baculovirus-infected silkworm larvae

**DOI:** 10.1101/2022.05.10.491283

**Authors:** Hiroyuki Hikida, Susumu Katsuma

## Abstract

*Bombyx mori nucleopolyhedrovirus* (BmNPV) is a pathogen of the domestic silkworm, *Bombyx mori*, whose infection drastically alters larval locomotory patterns and activity. As *B. mori* is the only insect that has been completely domesticated, its larvae are typically inactive. In contrast, BmNPV-infected larvae exhibit vigorous movement. Recently, we developed a method for locomotory analysis of BmNPV-infected *B. mori* larvae, which yielded high-resolution data on BmNPV-induced host locomotory alteration. However, this method had a disadvantage when analyzing the behavior of uninfected larvae, as they were typically inactive and difficult to distinguish from the background. In addition, the analysis processes involve many manual steps, which reduces analysis throughput. We have developed KaicoTracker, a new tool for automated locomotory analysis of BmNPV-infected *B. mori* larvae, using Python and OpenCV. This tool is an all-inclusive package that tracks and quantifies larval locomotion and verifies the accuracy of the tracking. Larval locomotion is tracked by motion tracking function based on k-Nearest Neighbor implemented in OpenCV. Based on the tracking results, the positions of the larvae are determined, and locomotory distance, speed, duration, and frequency are calculated. The larval positions are marked on the original video, allowing tracking precision to be examined. By analyzing previously obtained data, we confirmed that this tool provides a robust analysis of inactive larvae with fewer manual steps.

## Introduction

Baculovirus-induced host behavioral manipulation has been known for a long time in lepidopteran larvae. Infected larvae exhibit enhanced locomotory activity (ELA) and climbing behavior (CB), where larval horizontal and vertical locomotion, respectively, are altered (Vasconcelos *et al*., 1996; Goulson, 1997). Most ELA-inducing lepidopteran baculoviruses degrade larval bodies with viral cathepsin and chitinase at a late stage of infection, thereby releasing virions into the environment (Ohkawa *et al*., 1994; Hawtin *et al*., 1997). The combination of ELA and CB facilitates larval dispersal and the spread of contamination with the released virions. Also, these host behavioral manipulations result in host death at elevated positions, which is believed to facilitate spread of virions by rainfall and avian feeding (Entwistle *et al*., 1993; D’Amico and Elkinton, 1995).

*Bombyx mori nucleopolyhedrovirus* (BmNPV) is a member of baculovirus and a major pathogen of the domestic silkworm, *Bombyx mori. B. mori* is the only insect that has been completely domesticated, and its larvae are typically inactive. In contrast, BmNPV infection results in the larvae exhibiting distinct ELA (Kamita *et al*., 2005). In previous research, the BmNPV-induced ELA was quantified following the method developed for *Mamestra brassicae* larvae, in which larval locomotion was manually measured at multiple time points (Vasconcelos *et al*., 1996; Goulson, 1997). This conventional method was adequate to determine whether or not larval locomotion is upregulated, thereby identifying the genes involved in BmNPV-induced ELA (Kamita *et al*., 2005; Katsuma *et al*., 2012; Kokusho *et al*., 2015). Due to the limited resolution of the data, it was difficult to determine which locomotory properties (e.g., speed, duration, and frequency) were altered by the infection.

Recently, we developed a method that combines continuous observation with a time-lapse camera and computational image processing with ImageJ software (Hikida *et al*., 2020; Hikida and Katsuma, 2021). This method yielded high-resolution data, but had a disadvantage when detecting uninfected larvae. This method detects moving objects, based on a single background image calculated as the average of a large number of frames. However, as uninfected larvae are inactive, they are sometimes counted as background, resulting in substantial background noise in some instances. This drawback appears to be a significant issue for future applications of the method to analyze the phenotypes of mutant BmNPVs that fail to induce ELA. In addition, the method requires many manual steps, which reduces analysis throughput. In the present study, we sought to address these issues by developing a robust and automated tool for locomotory analysis of BmNPV-infected *B. mori*.

## Materials and Methods

### Analyzed dataset

Previous video data were used for analysis (Hikida and Katsuma, 2021). The data was divided into three videos, and frames from #234 to #29032, containing 24-h data from 72 to 96 hours post infection (hpi) were analyzed. The T3 strain (Maeda *et al*., 1985) was used as the wild-type BmNPV in this dataset.

### Preliminary larval detection

The background was estimated and subtracted from the input videos using k-Nearest Neighbor (KNN) algorithm (Zivkovic and Van Der Heijden, 2006), implemented in OpenCV with a learning rate of 0.8. Small noises were removed from background-subtracted images by averaging the values of each pixel (px) in a 10 × 10 px square. Each object’s contours were extracted from blurred images, and then approximated with circles. The X- and Y-coordinates of centers of circles were stored as the positions of moving objects. Since KNN requires multiple frames to construct stable background models, estimation of background models was started 100 frames before the first frame of analysis.

### Determination of analyzed area

The positions of moving objects were detected as described above, in the first 1000 frames of the input videos, and their kernel density estimates (KDEs) were calculated along the X- and Y-axes. Frames were subdivided into blocks at local minimums of the KDEs. A designated number of blocks were cropped for subsequent analysis by removing the peripheral blocks.

### Tracking of larval locomotion

Local minimums of KDEs were calculated from the table containing the X- and Y-coordinates of all the moving objects. Each frame was cropped and divided into blocks as described above. The mean X- and Y-coordinates of each block in a frame were calculated and used as the larval position in each frame. When no object was detected at the frame, a position in the previous frame was used as the current position.

### Quantification of larval locomotion

The larval locomotory distance at each time point was calculated as the distance between larval positions in two consecutive frames in each block. The distance was interpreted as the larval locomotory speed when the values were greater than 0 px, considering that the larva was moving at the time point. Otherwise, the larva was considered to be in a paused state. The sum of locomotory distances was presented as the total travel distance during the period of analysis.

The definition of continuous larval locomotion is as follows: When both the mean and 90% of larval locomotory distances in a set of consecutive frames were greater than 1 px, the larva was considered to be moving during the period. First, 11 sequential larval locomotory frames were examined to determine if the condition mentioned above was met. When the condition was met, a subsequent frame was added and tested. This process was repeated until the condition was no longer met; at that point the larva was considered to have stopped moving. The duration and frequency of each continuous locomotion over the period of analysis were calculated.

### Verification of larval tracking

The larval positions determined as described above were marked on the original videos. Videos were cropped and trimmed to contain the analyzed region and time frame, respectively.

### Statistical analysis

The rank-sum test was used to compare the total travel distance, median locomotory speed and duration, and locomotory frequency of mock- and T3-infected larvae (n = 6).

### Code availability

The source code and other usage details of the tool are available on GitHub (https://github.com/H-Hikida/KaicoTracker).

## Results and Discussion

### Workflow of KaicoTracker

Our previous method used Image J and Python for image and data processing, respectively. KaicoTracker, on the other hand, is an all-inclusive package based on OpenCV implemented in Python. The tool sequentially processes the input videos, returns figures and tables for larval locomotion, and verifies the accuracy of larval tracking (Fig. 1A). In image processing steps, moving objects in the original videos are detected, and their contours are depicted, which are subsequently approximated by circles (Figs. 1B–D). Larval positions are calculated based on the distribution of circles, and their accuracy is verified by marking the positions onto the original videos (Fig. 1E). In order to reduce memory usage, the distribution of objects is initially analyzed along the X- and Y-axes, and peripheral areas containing few objects are removed from the subsequent analysis (Fig. 1F). We implemented additional options so that users can tailor the parameters to their experimental conditions.

**Fig. 1.**
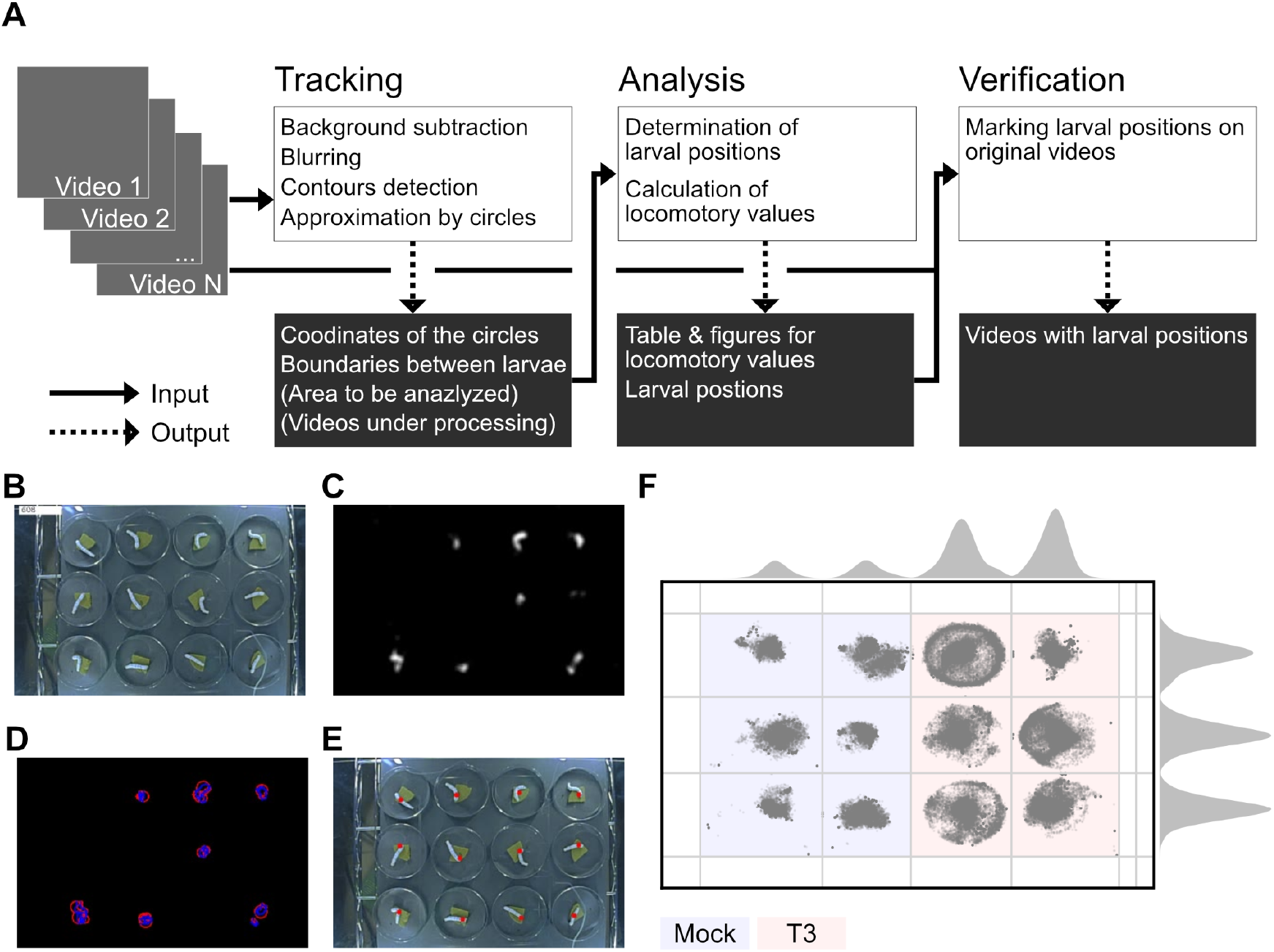
(A) A schematic workflow of KaicoTracker. Solid and dashed lines represent input and output, respectively. (B) An example frame from the input video along with its index. (C) A background-subtracted and blurred frame from (B). (D) Object detection from (C). Blue and red lines represent contours and circles approximating contours, respectively. (E) A frame indicating larval positions. Red dots represent larval positions. (F) A schematic illustration of boundary determination. Rectangle surrounded by a solid black line corresponds to the frame size of the video, while the gray dots represent larval positions. Colored backgrounds indicate the analyzed area, and blue and red indicate that for mock- and T3-infected larvae, respectively (n = 6). Gray lines indicate the boundaries between larvae. The plots at the top and right represent kernel density estimates for the X- and Y-axis distribution of the positions of moving objects, respectively.

### KNN-based motion tracking

The algorithm for detecting moving larvae is a major modification from the previous method in KaicoTracker. Previously, a single background image was constructed as an average of thousands of frames. This method intended to detect all larvae regardless of their locomotory state, but it incorrectly counted some larvae as background when they are in paused state for an extended period. It occasionally resulted in substantial noise, and our previous study failed to examine the locomotion of some uninfected larvae (Hikida and Katsuma, 2021). In contrast, KaicoTracker uses one of the machine-learning algorithms, KNN, implemented in OpenCV. KNN builds continuously updated models, estimated from several preceding frames, to separate images into foreground and background (Zivkovic and Van Der Heijden, 2006). The algorithm detects only moving larvae and disregards pausing ones (Fig. 1C). Therefore, long-term pausing of uninfected larvae has little effect on the detection accuracy.

### Reduction of manual steps

Another modification is the reduction of manual procedures. Previous methods required the manual determination of multiple parameters. One of the parameters is the size threshold for ignoring small noises. In KaicoTracker, foreground images were blurred, and these noises were automatically removed; therefore, a size threshold is not required (Fig. 1C). Another parameter is a boundary between larvae. In our experimental system, each larva was placed separately, and its locomotion was analyzed based on this separation. Previously, we determined the boundaries manually, but KaicoTracker automates this process using KDE (Fig. 1F). Larvae distributed in the center more frequently than on the periphery. When the distribution is smoothed by KDE, the X- and Y-coordinates display a clear peak around the center, and local minimums represent the boundaries. In addition, we implemented an option to calculate KDE from the first 1,000 frames, from which the edges of the analyzed area are determined, and frames are cropped at the edges (Fig. 1F).

### Example outputs

We applied KaicoTracker to analyze the previously collected data on the behavior of mock- and T3-infected *B. mori* larvae from 72 to 96 hpi (Hikida and Katsuma, 2021). We confirmed that mock-infected larvae periodically exhibited movement and stationary behavior, whereas T3-infected larvae lost this periodic pattern (Fig. 2A). The locomotory pattern was also disrupted, as T3-infected larvae moved to the peripheral area of the dish during later stages of observation, whereas mock-infected larvae remained in the center of the dish throughout the observation (Fig. 2B). These results are coincided with those obtained by the previous method (Hikida and Katsuma, 2021).

**Fig. 2.**
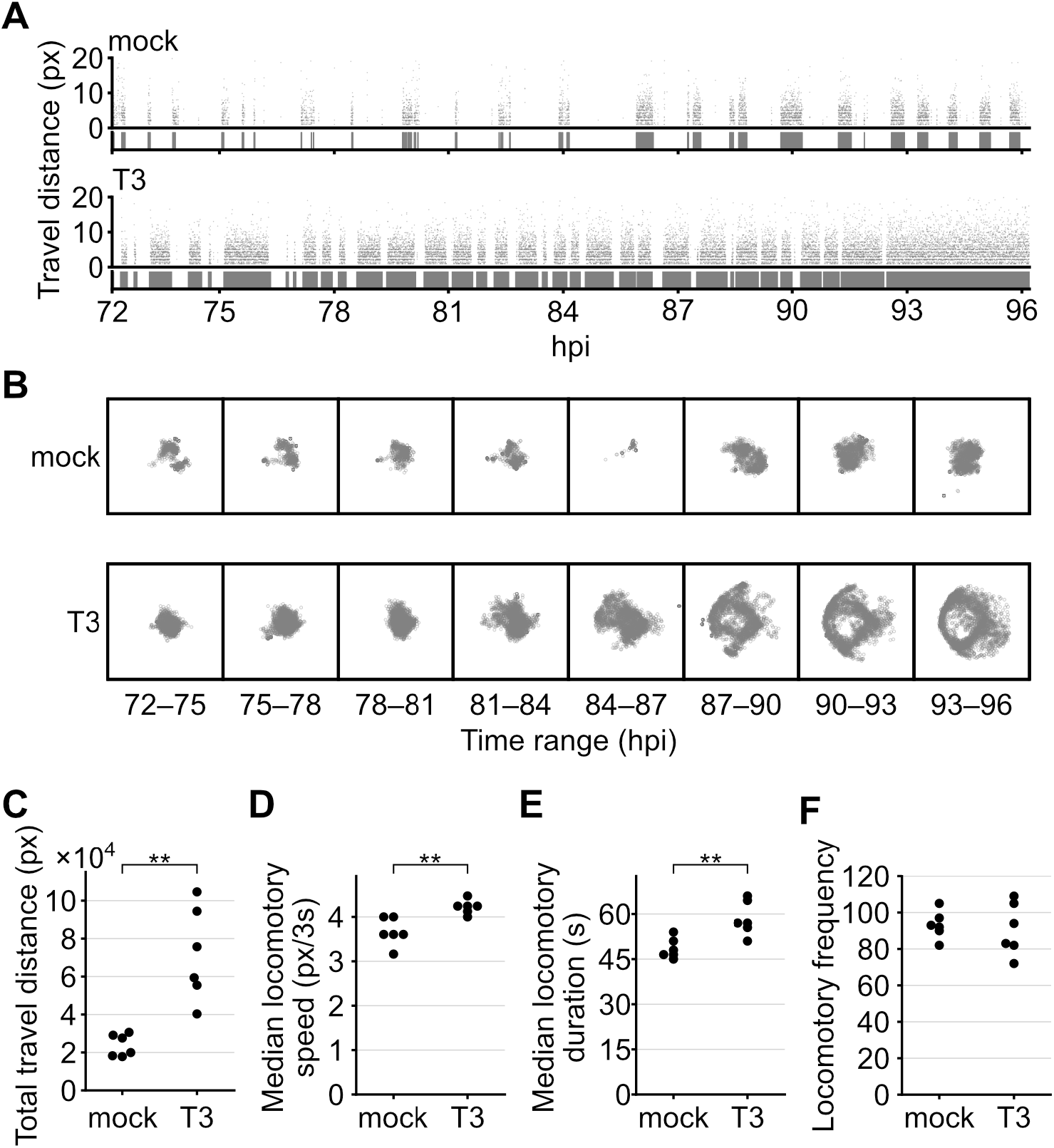
(A) The scatter plots and diagrams at the top and bottom represent locomotory distance at 3-s intervals and continuous locomotion, respectively. (B) Larval distribution for each 3-h interval. (A, B) One representative larva from mock- and T3-infected larvae, respectively. (C) The total travel distance in 24 h. (D) Median locomotory speed. (E) Median locomotory duration. (F) Locomotory frequency over 24-h period. (D–F) Dots indicate values determined for each larva. ** *p* < 0.01 by rank-sum test.

T3-infected larvae exhibited significantly greater total travel distance, higher locomotory speed, and longer locomotory duration, but similar locomotory frequency, compared with mock-infected larvae (Figs. 2C–F). In the previous report (Hikida and Katsuma, 2021), we were unable to calculate the locomotory values of a mock-infected larva due to the presence of substantial noise. In contrast, the present method enabled us to yield reasonable values for all mock-infected larvae using the same dataset. The result demonstrates that the newly developed method is robust for analyzing inactive larvae.

We found that absolute values varied from the previous report (Hikida and Katsuma, 2021). Compared to the previous values, the locomotory distance was higher, while the duration shorter. The present study employed different algorithms for tracking larval movement and calculating duration, which may have resulted in the differences of values. As shown in Fig. 1E, the larval positions were distributed in different parts of the larval bodies, and the detected pattern may vary between methods. Nevertheless, statistical analysis yielded the results consistent with those of the previous study (Hikida and Katsuma, 2021). Accurately highlighting the difference in larval locomotory values is a primary objective of both the previous and present methods. Therefore, we conclude that the current resolution is sufficient for quantitative comparisons.

## Conclusion

Here, we presented a novel tool for analyzing the locomotion of BmNPV-infected *B. mori* larvae. According to a reanalysis of the previous data, the newly applied algorithm is robust for analyzing inactive larvae. It indicates that this tool can be used to phenotype viruses with locomotory mutation. In addition, the majority of manual steps required in the previous method have been automated. The tool will facilitate a high-resolution locomotory analysis of BmNPV-infected *B. mori* and provide new insight into the underlying mechanisms.

## Acknowledgements

This study was supported by JSPS KAKENHI grant number 19J13438 to HH. HH was a JSPS Research Fellow (DC2).

